# Capsule and PspA Cooperatively Confer Resistance of *Streptococcus pneumoniae* to the Human defensin HNP-1

**DOI:** 10.1101/2025.09.29.679143

**Authors:** Thalita Bastos de Freitas e Silva, Maria Eduarda Pereira Mendes, Rebeca Faria, Kelvin Gattinoni, Bruna Terribile, Giulia Destro, Lucio F.C. Ferraz, Carlos J. Orihuela, Thiago R. Converso, Michelle Darrieux

**Affiliations:** Laboratório de Microbiologia Molecular e Clínica, Universidade São Francisco, Bragança Paulista, São Paulo, Brazil; Department of Microbiology, Heersink School of Medicine, The University of Alabama at Birmingham, Birmingham, Alabama, USA

**Keywords:** Pneumococcus, AMPs, HNP, defensin, PspA.6

## Abstract

*Streptococcus pneumoniae* resists host defenses through multiple virulence factors, yet their combined influence on the action of antimicrobial peptides remains unclear. We examined the role of Pneumococcal surface protein A (PspA) and the polysaccharide capsule in modulating susceptibility to the human defensin HNP-1. PspA-deficient strains of two different genetic backgrounds displayed increased sensitivity, while recombinant PspA neutralized peptide activity and anti-PspA antibodies enhanced bacterial killing. The capsule conferred serotype-dependent protection, with type 2 being more effective than type 4, and free polysaccharides acted as decoys by sequestering HNP-1. Removal of surface PspA from capsule-deficient mutants revealed additive contributions of both factors to survival. These findings highlight the complementary roles of capsule and PspA in pneumococcal resistance to HNP-1 and suggest that targeting these mechanisms could potentiate innate immune clearance and provide novel insights that may inform future vaccine design and antimicrobial strategies.

**AUTHOR SUMMARY:** *Streptococcus pneumoniae* causes serious infections such as pneumonia and meningitis, in part by evading the human immune system. One key component of our immune defense is antimicrobial peptides like HNP-1, which directly kill bacteria. In this study, we investigated how two major pneumococcal virulence factors – the surface protein PspA and the sugar capsule – help the bacterium resist killing by HNP-1. We found that removing PspA made the bacteria more susceptible, while adding purified PspA or blocking it with antibodies increased the peptide activity. The protective effect of the capsule depended on the sugar composition, and purified capsule sugars could bind and neutralize HNP-1, limiting its activity against pneumococci. When we removed PspA from bacteria lacking a capsule, the bacteria became even more sensitive, showing that both factors contribute to resisting immune attack. These results reveal how *S. pneumoniae* uses multiple strategies to survive innate immune defenses and highlight potential targets for improved vaccines or new treatments.

## 1. INTRODUCTION

*Streptococcus pneumoniae* remains a major global pathogen responsible for localized and systemic infections, such as otitis media, sinusitis, pneumonia, and meningitis (1). Its pathogenic success is largely due to its ability to asymptomatically colonize the nasopharyngeal mucosa, from where it can disseminate and cause disease (2). An important factor contributing to pneumococcal dispersion and spread from the upper respiratory tract is coinfection with viruses, particularly, influenza, which damages the mucosal barrier and triggers genetic changes in the bacteria resulting in a more invasive phenotype (2, 3). *S. pneumoniae* is among the leading causes of community-acquired pneumonia in many countries, affecting mainly children under five years of age and older adults, and remains one of the most important causes of bacterial meningitis. Pneumococcal infections are responsible for more than one million deaths worldwide each year, principally among children and elderly (4).

The polysaccharide capsule is an essential virulence factor in *S. pneumoniae*; its principal role is to act as a barrier that inhibits phagocytosis by the host’s immune cells (5, 6). Capsule also prevents antibody binding to surface proteins (7), promotes colonization of the nasopharynx (8) and confers resistance to intracellular killing by vascular endothelial cells during invasive disease (9). The absence of capsule has been associated with an almost absolute reduction in the ability to cause invasive infection; therefore, capsular polysaccharides have been used as the basis for pneumococcal vaccines, such as the 23-valent polysaccharide vaccine (PPV23) and conjugate vaccines (PCV), which have demonstrated considerable efficacy in reducing the incidence of pneumoniae and invasive disease (10, 11). Critically and despite being immunogenic in immunocompetent individuals, capsular polysaccharides display great structural and serological diversity, with more than 100 distinct serotypes identified so far (12).

Another important virulence factor in pneumococcus is Pneumococcal surface protein A (PspA), an exposed protein noncovalently linked to phosphorylcholine residues which compose part of the bacterium’s lipoteichoic and cell wall associated teichoic acid. PspA displays multifunctional roles including inhibition of Complement System activation/deposition on the bacterial surface, interaction with host molecules such as glyceraldehyde-3-phosphate dehydrogenase (GAPDH) and lactate dehydrogenase (LDH), co-opting of GAPDH and LDH released by dying host cells for the bacterium’s benefit (13, 14), and interference with antimicrobial peptides (AMPs) (15). Recent studies also suggest that PspA may play a role in membrane remodeling, an essential mechanism for maintaining cellular functionality in response to stress (16). The importance of PspA is best evidenced by the fact that mutants deficient in PspA are highly attenuated *in vivo* (17–19). Notably, the structure of PspA includes three main regions: a variable N-terminal portion that protrudes from the capsule and is accessible to the host’s immune system, a proline-rich region which connects the N- and C-termini and includes protective epitopes (20) and a more conserved C-terminal region, responsible for anchoring the protein to the bacterial surface. Based on variations in a region within the N-terminal domain called clade-defining region (CDR), PspA has been classified in three families and six clades (21). Variation in the host-factor binding domains within the N-terminal also means that PspA has different roles in different strains (13, 14).

Based on its exposure on the bacterial surface, prevalence among clinical isolates and important role in pathogenesis, PspA has been evaluated as a potential serotype independent vaccine against pneumococcal infections, with encouraging results (22, 23). It is highly immunogenic, inducing specific antibodies both in convalescent patients and in experimental models, and protective against pneumonia and invasive disease in different animal models.

The interactions between pneumococcus and the immune system are also influenced by antimicrobial peptides (AMPs), which are critical components of the innate response. Human defensins, particularly human neutrophil peptides (HNPs), such as HNP-1, are secreted by neutrophils and possess antiviral and antibacterial activities (24). Research indicates that the interaction between cationic AMPs and the negatively charged bacterial surface results in depolarization and pore formation, leading to membrane disruption (24, 25). In addition to this direct membrane permeabilization, defensins can interfere with cell wall synthesis through interaction with Lipid II (26). Human α-defensins have also been shown to neutralize pneumococcal virulence by inhibiting cholesterol-dependent cytolysins, thereby reducing toxin-mediated cell damage (27). However, the pneumococcus has developed effective strategies to resist these AMPs, including the use of efflux pumps, sequestration of antimicrobial molecules, and modifications to the cell wall that alter the capsule structure (28).

While the individual contributions of the polysaccharide capsule and PspA to pneumococcal resistance against some antimicrobial peptides (AMPs) have been established (29–32), a critical knowledge gap remains regarding their combined or potentially additive role against human defensins. Notably, the role of the capsule in pneumococcal resistance to HNP-1 has yielded conflicting findings, with some studies reporting a protective effect and others indicating increased susceptibility (33, 34), and the specific contribution of PspA on the activity of defensins has yet to be elucidated. Therefore, the present study investigates the respective and combined contributions of PspA and the polysaccharide capsule to pneumococcal resistance against the human neutrophil peptide HNP-1.

## 2. RESULTS

### 2.1 PspA contributes to resistance against HNP-1

To assess the role of PspA in resistance to HNP-1, wild-type strains and their corresponding PspA-negative mutants were exposed to different concentrations of HNP-1 for one hour, and bacterial viability was determined by plate count. Figure 1 shows the percent survival of the wild-type strains of serotypes 2 and 4, in comparison with their PspA-negative isogenic mutants, after treatment. In both strains, the absence of PspA resulted in significantly lower resistance to all concentrations of HNP-1, in a dose-dependent effect For serotype 2, strain D39 (expressing Family 1, clade 2 PspA) the wild-type version was highly resistant to lower peptide concentrations of 3 and 6.25 µg/ml (Figure 1A), while the PspA deficient mutant presented significant reduction in viability, with 30 to 40% killing. At the dose of 12.5 µg/ml of peptide, mean survival was 50% in the mutant, against 70% in the parental strain. The highest concentration of HNP-1 resulted in the greatest difference in survival, reducing the PspA-negative strain to less than half the survival of the wildtype strain. For serotype 4, using the TIGR4 strain (expressing Family 2 PspA) and its PspA-negative mutant BR61.1, the effect of the defensin was considerably greater at lower concentrations. Moreover, the bacterium overall was more sensitive to killing. The mutant exhibited a marked reduction in viability starting from the lowest AMP concentration (3 µg/ml), with only 20% survival. In contrast, its wild-type counterpart reached 50% bacterial killing only at a concentration of 12.5 µg/ml (Figure 1b). At the higher concentration of the HNP-1, the mutant strain was completely abrogated, while the parental strain still showed 20% survival. Taken together, these findings indicate that PspA is important for protection against HNP-1, and this varies between strains, however other factors are also important.

**Figure 1.**
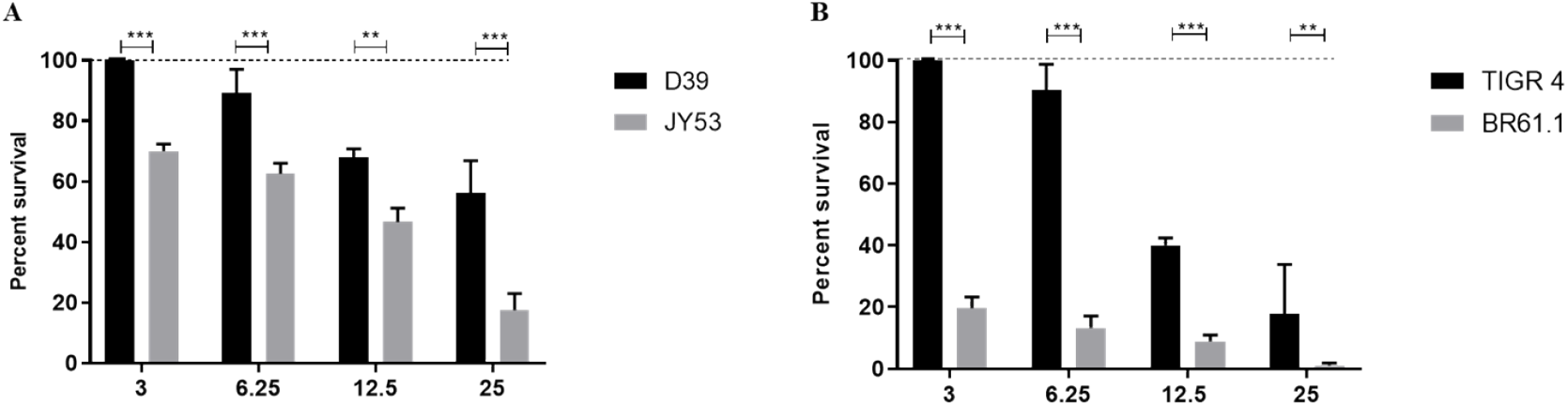
PspA expression affects *Streptococcus pneumoniae* resistance to HNP-1. The bacteria were treated with increasing concentrations of the AMP (3, 6.25, 12.5 and 25 µg/ml and survival were compared among groups using two-way ANOVA with Sidak post-test. Data represents two independent experiments; each performed with four replicates per group. Each bar represents the percent survival in relation to untreated controls (dashed lines). A) Comparison between wild type serotype 2 strain (D39) and its isogenic PspA mutant, JY53. B) Comparison between wild type serotype 4 strain (TIGR4) and its isogenic PspA mutant, BR61.1. *p<0,05; ** p<0,01; ***p<0,001 when comparing different bacteria treated with the same concentration of HNP-1.

### 2.2 Recombinant PspA neutralizes the bactericidal activity of HNP-1

Given the increased susceptibility of PspA-negative strains to HNP-1, we investigated whether free recombinant PspA (rPspA) could neutralize its observed bactericidal effect. To test this, HNP-1 (25 µg/ml) was pre-incubated with rPspA fragments corresponding to strain P94 also family 1, clade 2, before being added to the D39 strain. As shown in Figure 2, the addition of rPspA led to a modest but significant increase in bacterial survival, compared to the control treated with HNP-1 alone (Figure 2). These results demonstrate the protective effect of bacteria-free PspA, against HNP-1-mediated killing.

**Figure 2.**
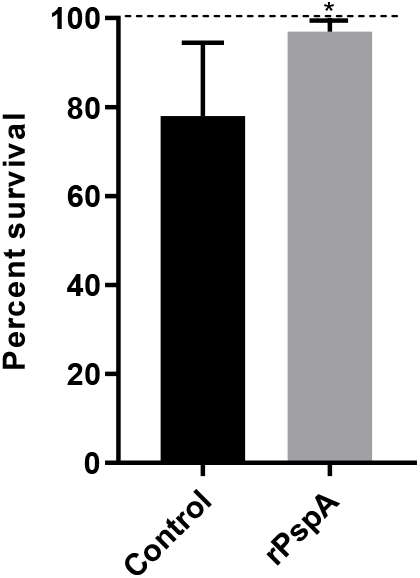
Exogenous PspA limits *Streptococcus pneumoniae* killing by HNP-1. *Streptococcus pneumoniae* D39 was treated with 25 µg/ml of HNP-1 previously incubated with a recombinant N-terminal PspA fragment (rPspA). The control group was incubated with HNP-1 and BSA. Data represents two independent experiments; each performed with four replicates per group. Each bar represents percent survival in comparison with untreated bacteria (dashed line). Statistical analysis was performed using Student t test. *p<0,05 in comparison with the control.

### 2.3 Anti-PspA antibodies sensitize *S. pneumoniae* to killing by HNP-1

To assess the influence of anti-PspA antibodies on the bactericidal activity of HNP-1, D39 were opsonized with sera from mice previously immunized with recombinant PspA from strain P94 prior to peptide exposure. This resulted in significantly enhanced bacterial killing compared to the control group opsonized with control serum and treated with HNP-1, as shown in Figure 3. In context of the more modest protection seen with exogenous added PspA (Fig 2), this finding suggests that surface bound PspA is more effective in protecting the bacterium from HNP-1.

**Figure 3.**
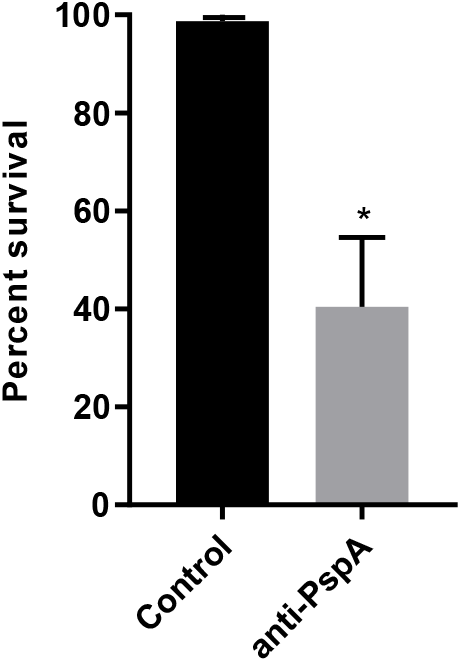
Anti-PspA antibodies enhance *Streptococcus pneumoniae* killing by HNP-1. The peptide was added to *Streptococcus pneumoniae* D39 previously opsonized with 5% sera from mice immunized with rPspA, at a concentration of 25 µg/ml. The control group was incubated with sera from mice injected with adjuvant diluted in saline, and HNP-1. Data represents two independent experiments; each performed with four replicates per group. Each bar represents percent survival in comparison with the control. Statistical analysis was performed using Student t test. *p<0,05 in comparison with control serum.

### 2.4 The role of the capsule in pneumococcal resistance to HNP-1 is serotype-dependent

D39 and TIGR4 have distinct capsular types and we sought to determine the contribution of the polysaccharide capsule in HNP-1 resistance. This was done by treating wild-type strains and their isogenic capsule-negative mutants with increasing concentrations of the AMP. A protective role for the serotype 2 capsule was observed when comparing the wild-type D39 strain to its nonencapsulated mutant, AM1000 (Figure 4a). Susceptibility of the AM1000 mutant was evident starting at a concentration of 6.25 µg/ml, with a 100% inhibition of bacterial survival at the highest concentrations (12.5 and 25 µg/ml). In contrast, the encapsulated wild-type strain showed a maximum 40% reduction in viability, even at the highest tested concentration.

**Figure 4.**
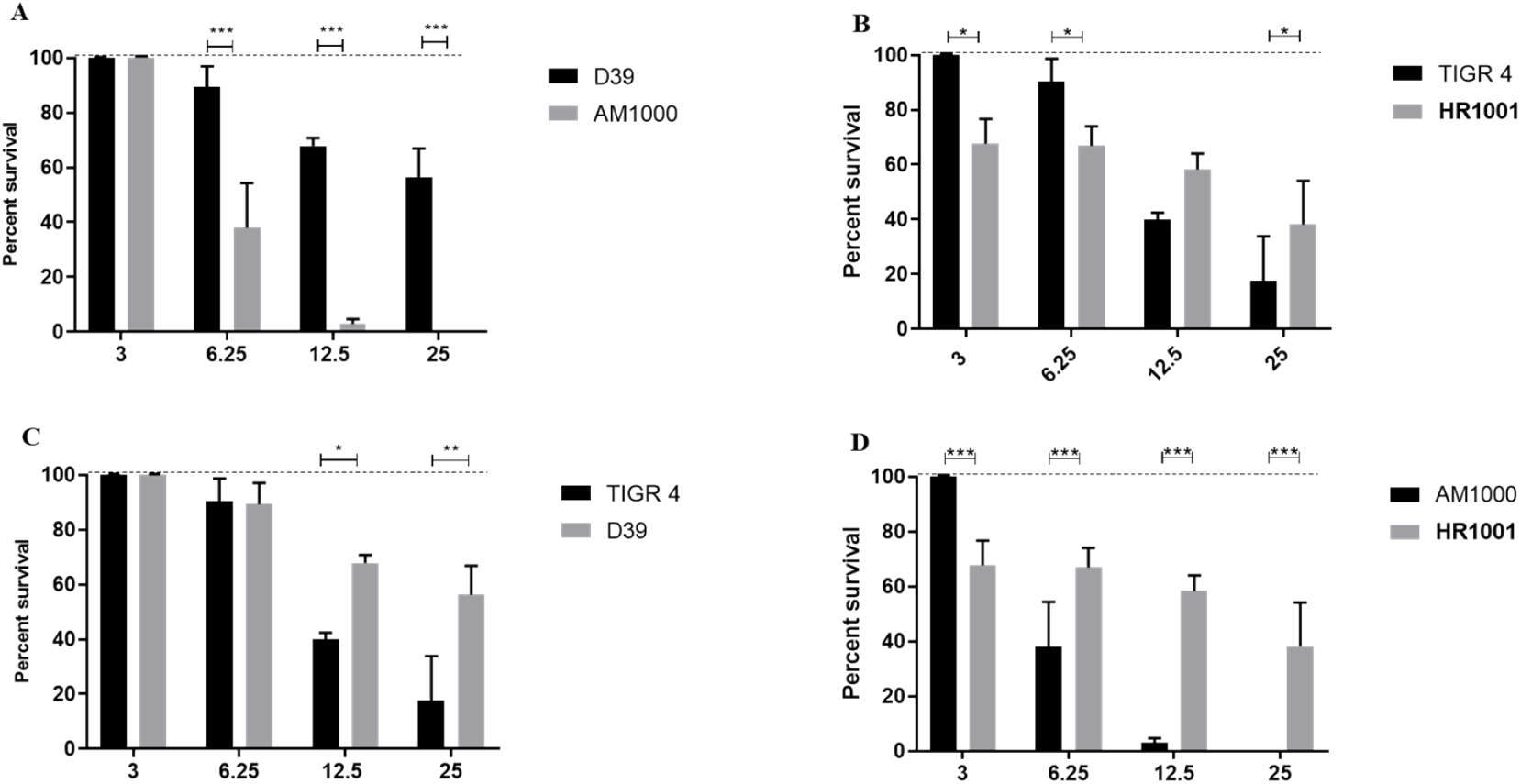
Effects of capsule on *Streptococcus pneumoniae* resistance to HNP-1. The bacteria were treated with increasing concentrations of the AMP (3, 6.25, 12.5 and 25 µg/ml) and survival was compared among groups using two-way ANOVA with Sidak post-test. Data represents two independent experiments; each performed with four replicates per group. Each bar represents the percent survival in relation to untreated controls (dashed lines). A) Comparison between wild type serotype 2 strain (D39) and its isogenic capsule negative mutant, AM1000. B) Comparison between wild type serotype 4 strain (TIGR4) and its isogenic capsule negative mutant, HR1001. C) Comparison between two pneumococcal serotypes, 2 (D39) and 4 (TIGR4). D) Comparison between the two capsule-negative mutants, AM1000 and HR1001. *p<0,05; ** p<0,01; ***p<0,001 when comparing different bacteria treated with the same concentration of HNP-1.

On the other hand, the effects of HNP-1 treatment in the serotype 4 strain TIGR4 and its isogenic capsule-deficient mutant, HR1001 varied depending on the peptide concentration. The parental strain was more resistant than the mutant in the lower concentrations of HNP-1 (3 and 6.25 µg/ml), while it tended to become more susceptible as the HNP-1 concentration increased. In the highest concentration of the peptide (25 µg/ml), the mutant strain was significantly more resistant to the AMP.

A comparison between the wild-type strains, D39 (serotype 2) and TIGR4 (serotype 4), revealed marked differences in susceptibility to the peptide (Figure 4c). Although both strains were resistant to the lower concentrations of HNP-1, the D39 strain was significantly more resistant than TIGR4 in AMP concentrations of 12.5 and 25 µg/ml. The capsule mutant strains also differed in their susceptibility to killing by HNP-I (Figure 4D). The capsule-4 derivative, HR1001, was significantly more resistant to AMP concentrations of 6.25 µg/ml and higher than AM1000. When compared with the results from the PspA-negative strains, it is possible to infer that in strain D39, capsule has a more pronounced effect of resistance to HNP-1 than does PspA, whereas in strain TIGR4, the reverse is apparent. This result suggests that the effects of capsule on pneumococcal resistance to HNP-1 vary among different genetic backgrounds and this is likely due in part to differences in capsule type.

### 2.5 Free capsular polysaccharides partially neutralize the bactericidal activity of HNP-1

To determine whether free capsular polysaccharide could neutralize the bactericidal action of HNP-1, the peptide was pre-incubated with either serotype 2 (PS-2) or serotype 4 (PS-4) polysaccharide prior to bacterial exposure. As shown in Figure 5A, HNP-1 pre-treated with free PS led to a significant increase in D39 survival when compared to treatment with HNP-1 alone: the group incubated with PS-2 showed around 90% survival, against less than 60% survival after treatment with the peptide alone. A more moderate effect was observed for PS-4 plus HNP-1 treatment in TIGR4 (figure 5D): HNP-1 alone reduced bacterial viability to less than 20%, while HNP-1 combined with PS-4 showed a slight increase in survival, to around 25%.

**Figure 5.**
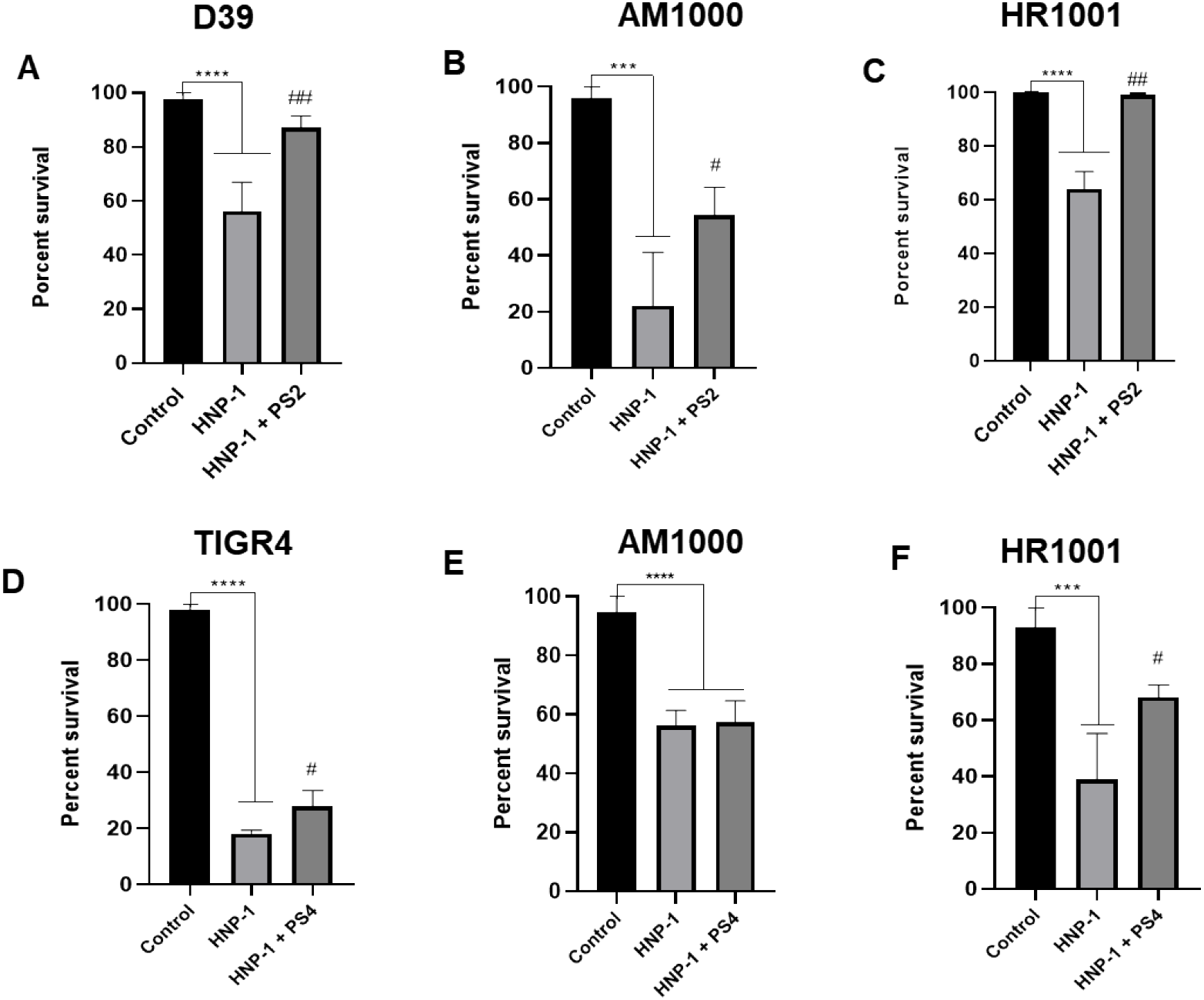
Effects of free capsular polysaccharide in *Streptococcus pneumoniae* resistance to HNP-1. Wild type strains D39 and TIGR4, and its isogenic capsular-negative mutants AM1000 and HR1001 were treated with 25 µg/ml of HNP-1 alone or in combination with 10 µg/ml of purified PS-2 (A-C) or PS-4 (D-F). Data represents two independent experiments; each performed with four replicates per group. Each bar represents the percentage of bacteria surviving treatment, in comparison with the untreated control. Comparison among groups was performed using one-way ANOVA with Tuckey post-test. ** p<0,01 in comparison with untreated control. #p<0,05 when comparing treatment with HNP-1 alone and in presence of purified PS-2 or PS-4.

The effect of free polysaccharide was also investigated in capsule-negative mutants, AM1000 and HR1001. Addition of free PS-2 significantly protected both capsule-negative mutants from HNP-1 induced killing (Figure 5B and C). On the other hand, the presence of PS-4 led to an increased survival only in the HR1001 mutant; the AM1000 strain showed similar killing with or without PS-4. Taken together, these results demonstrate that the effect of free-polysaccharides in pneumococcal resistance to HNP-I are dependent on their biochemical constitution, with PS-2 being more effective in neutralizing the peptide than PS-4.

### 2.6 PspA and capsule additively contribute to HNP-1 resistance

Finally, and to further investigate the combined effects of PspA and capsule in pneumococcal resistance to HNP-1, the capsule-negative mutants were washed with choline chloride to remove surface-bound PspA and then exposed to HNP-1. In the serotype 2-derived mutant, AM1000, removal of PspA led to a marked reduction in bacterial viability (Figure 6a), demonstrating a clear additive protective effect between PspA and the capsule for this strain.

**Figure 6.**
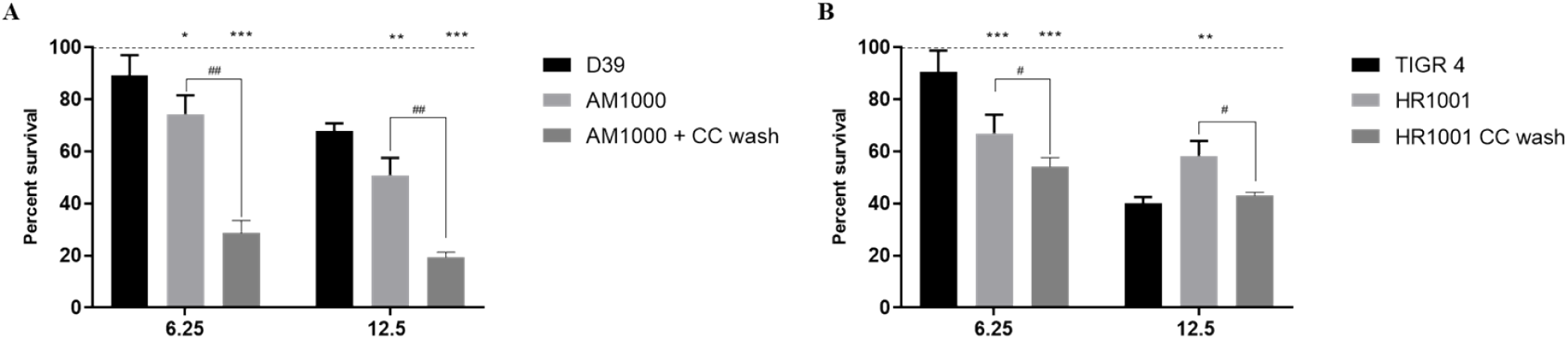
Combined effects of PspA and capsule on *Streptococcus pneumoniae* resistance to HNP-1. Two capsule-negative mutans, AM1000 (A) and HR1001 (B) were washed with choline-chloride to remove PspA and treated with two concentrations of the AMP (6.25 and 12.5 µg/ml) and survival was compared among capsule producing, capsule negative, and capsule negative + PspA removed groups Data represents two independent experiments, each performed with four replicates per group. Statistical analysis was performed using two-way ANOVA with Sidak post-test. Each bar represents the percent survival in relation to untreated controls (dashed lines). *p<0,05 in comparison with the parental strain; ***p<0,001 in comparison with the parental strain; ^##^p<0,001 when comparing intact and cc-washed bacteria.

In contrast, a more complex interaction was observed in the serotype 4-derived capsule negative strain, HR1001. Removal of PspA hindered the bacterium more susceptible to the lower concentration of the AMP, 6.25 µg/ml. However, when the AMP concentration was enhanced to 12.5 µg/ml, PspA removal did not aid in bacterial killing, presenting similar survival as the wild-type strain, TIGR4 (Figure 6b). Interestingly, there is a significant reduction in bacterial viability when comparing the capsule-negative derivative of TIGR4, HR1001, before and after PspA removal, indicating that the loss of PspA favors HNP-1 activity in this resistant non encapsulated mutant. Taken together, these results reveal that while PspA consistently confers protection, its combined effect with the polysaccharide capsule on HNP-1 resistance is dependent on the pneumococcal serotype.

## 3. DISCUSSION

The polysaccharide capsule and the surface protein PspA are recognized as central virulence factors in *Streptococcus pneumoniae*, acting independently and complementarily in immune evasion and resistance to antimicrobial molecules (6, 15, 46). Thus, the present study aimed to investigate the individual and combined contributions of PspA and capsule to the bactericidal action of the human defensin HNP-1.

A comparative analysis of wild type and PspA-negative mutant pneumococcal strains revealed an increased sensitivity of the mutants to treatment with HNP-1. This effect was independent of capsular type, as shown by the significantly reduced survival of PspA-negative mutants derived from both serotype 2 and serotype 4 strains. These results suggest that PspA plays an important role in protection against the antimicrobial peptide HNP-1, as its absence renders pneumococci more susceptible even in the presence of an intact capsule. Supporting these findings, we demonstrate that addition of a recombinant PspA fragment can rescue pneumococci from the bactericidal activity of the peptide, increasing survival upon HNP-1 treatment. This piece of evidence further reinforces PspA’s protective efficacy, while indicating that the protein does not need to be attached to the bacteria to exert its effect.

Previous work evaluating the interactions of pneumococci with antimicrobial peptides have found that PspA can protect the bacterium from apolactoferrin – the bactericidal form of the defense protein lactoferrin (29, 32, 47) and the bovine cathelicidin, indolicidin (30). In both cases, PspA has been shown to specifically bind to the antimicrobial proteins (30, 48). These studies demonstrate that PspA exhibits broad-spectrum AMP-sequestering activity, which, together with its well-documented ability to inhibit complement activation, contributes to pneumococcal survival during both colonization and invasive disease.

Since PspA has long been investigated as a potential vaccine candidate against pneumococcal disease, we sought to determine how anti-PspA antibodies affect HNP-1 mediated killing. Pre-opsonization with serum from mice which had been immunized with a recombinant N-terminal PspA fragment enhanced the bactericidal effect of HNP-1 in comparison with control serum, likely by inhibiting PspA’s capacity to bind the peptide, thereby allowing HNP-1 to access the bacterial membrane and exert its disruptive effect. These observations support the notion that antibodies induced by vaccination with recombinant N-terminal fragments of PspA may confer protection not only by promoting opsonophagocytosis (45), but also by unmasking the action of innate immune effectors such as antimicrobial peptides, thereby enhancing bacterial clearance at mucosal and systemic sites.

Regarding the capsule, we observed that serotype 2 (D39) showed greater resistance to HNP-1 than serotype 4 (TIGR4). Additionally, the loss of capsule had variable effects depending on the bacterial serotype: the capsule negative mutant derived from serotype 2 showed reduced survival in comparison with the parental strain, while the non-encapsulated mutant from TIGR4 was more sensitive to the lower concentrations of HNP and became more resistant as the AMP concentration increased. These results indicate that not merely the presence of capsule, but its composition has a profound impact on the pneumococcal ability to resist the bactericidal action of the human defensin HNP-1.

Insights into the possible mechanisms underlying this effect suggest that the surface electric charge may help shield the bacterium from cationic peptides such as HNP-1. Previous data by our research group has shown that denser capsules with a higher negative charge offer greater protection against indolicidin, a cationic peptide belonging to the cathelicidin family (31). Accordingly, the serotype 2 strain D39 has a much higher electronegativity than the capsule 4 strain, suggesting that the capsule charge may influence pneumococcal resistance to different cationic antimicrobial peptides. A similar protective effect of the polysaccharide capsule against AMPs has been demonstrated by (33). In that study, the addition of negatively charged pneumococcal polysaccharide, but not neutral or positive molecules, protected unencapsulated bacteria from killing by HNP-1. Moreover, addition of polycations restored AMP activity, indicating that the negative charge is essential for the protective effects of polysaccharides. The authors propose that the capsule may act as a “decoy” by attracting the peptides away from the membrane, thus preventing their bactericidal activity. Our findings, therefore, suggest that the effect of the capsule on the action of HNP-1 is dependent on both the serotype and the concentration of the peptide, which is in line with recent studies that highlight the structural diversity of capsular polysaccharides and their impact on colonization and immune response (49–51).

Contrasting those results, Beiter et al. (2008) (34) observed that non-encapsulated mutant pneumococci were more resistant to treatment with human defensins than their respective parental strains. This variation could be attributed to differences in the peptide: the authors used a mixture of HNP-1 to III purified from human neutrophils, instead of isolated HNP-1. It is also important to notice that in that study, the D39 strain did not show reduced resistance to the AMP, but rather a comparable survival to the capsule negative derivative.

Another relevant point was the observation that purified polysaccharides (PS-2 and PS-4) were able to partially neutralize the activity of HNP-1. This effect was more evident with PS-2, suggesting a serotype-specific role in modulating resistance. This mechanism is compatible with the molecular “decoy” hypothesis described by Llobet et al. (2008), in which bacterial capsules, even in free form, act by diverting antimicrobial peptides and reducing their efficacy (33). That study has also shown that AMP treatment promotes capsule release, resulting in an entrapment of the peptide to prevent killing. Further studies have confirmed that capsule shedding promoted by cationic AMPs in the mucosal sites is an important mechanism for pneumococcal colonization, allowing the exposure of adhesins at the bacterial surface that favor interaction with epithelial cells, while the free polysaccharides sequester the AMPs and prevent killing (52). The mechanism of capsule shedding is dependent on the cell wall hydrolytic activity of the suicidal amidase autolysin LytA, which promotes a restructuring of the bacterial surface and favors epithelial invasion (52).

To evaluate the additive contributions of capsule and PspA to pneumococcal resistance against HNP-1, two capsule-negative mutant strains were incubated with choline chloride to remove surface-attached PspA, prior to treatment with the peptide. The combined analysis revealed that capsule and PspA both play a role in protecting against HNP-1, especially in serotype 2, where the absence of capsule and PspA greatly impaired survival. In serotype 4, although the capsule mutant strain showed increased resistance to the higher concentration of HNP-1, PspA removal still significantly reduced survival, suggesting that resistance to HNP-1 results from a dynamic equilibrium between multiple surface factors. This variation may be associated with both biochemical differences of capsular polysaccharides and differential expression of cell wall-associated proteins, as described by Mitchell and Mitchell (2010) and Abdul Rahman et al. (2023) (5, 12).

Although our study functionally demonstrates the contribution of both capsule and PspA to HNP-1 resistance, the underlying molecular mechanisms remain speculative. Potential mechanisms that could prevent the peptide from reaching the bacterial membrane include the direct sequestration of HNP-1 by surface components or electrostatic repulsion mediated by surface charge. Therefore, future studies employing different techniques are necessary to clarify these binding interactions and provide a detailed mechanism basis for the observed resistance. Recognition of these mechanisms is essential for understanding the pathogenesis of pneumococcus and for developing alternative therapeutic strategies. In this sense, the combination of defensins with agents that inhibit capsular synthesis or block the action of PspA may represent a promising approach in the face of increasing antimicrobial resistance (53, 54). In conclusion, our results reinforce the multifaceted role of capsule and PspA in the resistance of *S. pneumoniae* against human defensins and indicate that future prevention and treatment strategies should consider not only serotype variability, but also the contribution of surface proteins in the modulation of host response.

## 4. MATERIALS AND METHODS

### 4.1 Bacterial strains and growth conditions

Table 1 summarizes the pneumococcal strains and mutants used in this study. The bacteria were kept as frozen stocks at −80º and, on the day anteceding the experiments, plated on blood agar (BA) and incubated for 18-20 h at 37ºC in microaerophilic conditions. The next day, the colonies were transferred to 5 mL of Todd Hewett broth supplemented with 0.5% yeast extract (THY) and cultured until they reached an optical density (O.D._600nm_) of 0.3-0.4 (approximately 10^7 CFU/mL). The mutant strains are part of a collection maintained at the University of Alabama at Birmingham (UAB) and kindly provided by Dr Anders Hakansson from Lund University (LU).

**Table 1.**
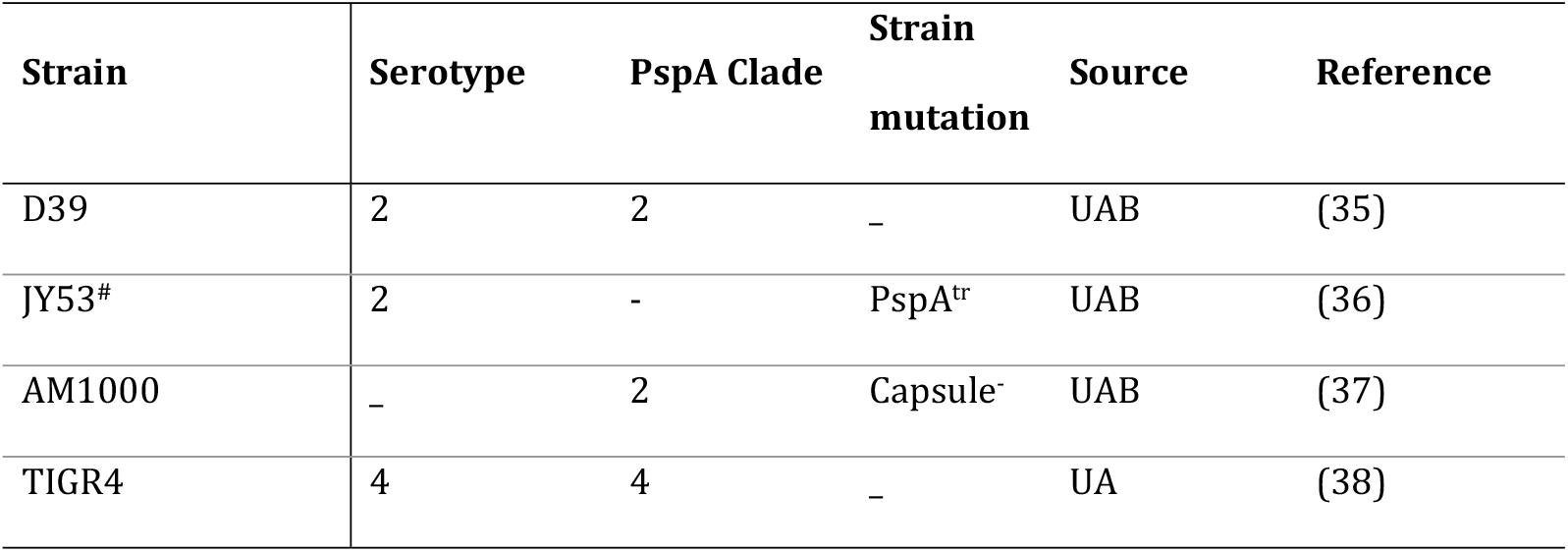
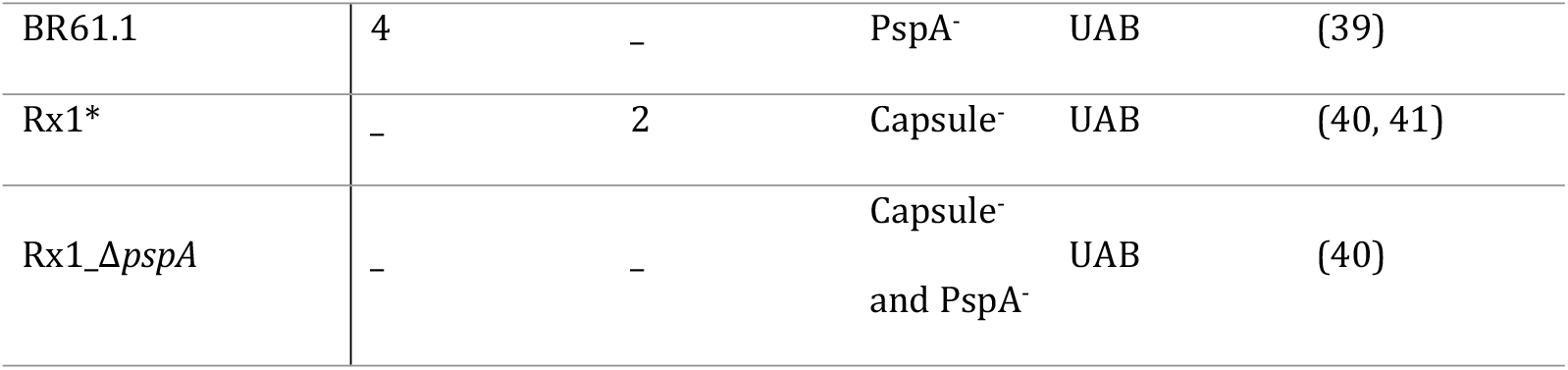
Pneumococcal strains used in this study. UAB – University of Alabama at Birmingham, USA.

| Strain | Serotype | PspA Clade | Strain mutation | Source | Reference |
| --- | --- | --- | --- | --- | --- |
| D39 | 2 | 2 | – | UAB | (35) |
| JY53<sup>#</sup> | 2 | - | PspA<sup>tr</sup> | UAB | (36) |
| AM1000 | – | 2 | Capsule<sup>-</sup> | UAB | (37) |
| TIGR4 | 4 | 4 | – | UA | (38) |
| BR61.1 | 4 | – | PspA<sup>–</sup> | UAB | (39) |
| Rx1* | – | 2 | Capsule<sup>–</sup> | UAB | (40, 41) |
| Rx1_Δ*pspA* | – | – | Capsule<sup>–</sup><br>and PspA<sup>–</sup> | UAB | (40) |

### 4.2 *In vitro* Bactericidal Assay

Pneumococcal susceptibility to HNP-1 was determined using *in vitro* bactericidal assay adapted by Waz et al 2024 (30). Once the cultures reached the desired O.D._600nm_, 2 ml of the culture were pelleted by centrifugation at 5.000 rpm for 5 minutes, washed with 1 ml of PBS and resuspended with an equal volume of sterile 10 mM sodium phosphate buffer at pH 7.6. Cationic adjustment was performed according to the protocol described by Sánches et al. and Yin et al. (42, 43) to achieve similar ionic levels to those found in human tissues. The bacterial suspension was then subjected to treatment with different concentrations (ranging from 3 to 25 µg/ml) of HNP-1 (ANASPEC code AS-60743), which was diluted in acetic acid + bovine serum albumin (BSA) and kept at −20ºC. At the time of use, HNP-1 was further diluted in sterile 10 mM sodium phosphate buffer at pH 7.6. The control group was incubated with phosphate buffer alone. After one hour incubation at 37ºC, the samples were subjected to serial dilution, plated on BA and incubated at 37 ºC for 16-20 h. The number of surviving bacteria after treatment was calculated and compared among the groups.

To assess the contribution of the polysaccharide capsule to HNP-1 bactericidal action on pneumococci, wild-type pneumococci and isogenic mutants that do not produce capsule, were treated with increasing concentrations of the peptide and plated as described. Similarly, the effect of PspA expression on pneumococcal susceptibility to HNP-1 was evaluated by comparing wild-type strains D39 and TIGR4 with their isogenic Δ*pspA* mutants (JY53 and BR61.1, respectively).

### 4.3 Recombinant PspAs

The PspA fragment used in this study was derived from strain P94 (family 1, clade 2) (44, 45). The gene fragments encompassing the sequences for the N-terminal alpha-helical region and the proline-rich domain (Supplementary Figure 1), were amplified from the chromosomal DNA and inserted into the pAE-6xHis vector. Protein expression was carried out in *E. coli* BL21DE3, followed by purification via affinity chromatography, following the method outlined by Converso et al., 2017 (45).

### 4.4 Production of anti-PspA and anti-capsular antibodies

The assays using serum from animals immunized with rPspA were conducted after approval by the Ethics Committee on Laboratory Animal Research at São Francisco University (CEUA/USF, protocol 001.06.2019). Animals were immunized via subcutaneous injection with 10 μg of rPspA in three consecutive doses at 10-day intervals, using Alum (Al(OH)_3_,50 μg/animal/dose) as an adjuvant. Control mice were injected with the adjuvant diluted in saline solution. After the final immunization, 100 μL of blood was collected through retro-orbital puncture after local anesthesia. Pooled sera from five mice immunized with the same protein was separated by centrifugation and stored at −20 ºC for subsequent analysis. Specific IgG levels were determined by ELISA.

### 4.5 Effect of rPspA and purified PS addition on HNP-1 activity

The effect of excess PspA and capsular polysaccharide on pneumococcal susceptibility to HNP-1 was evaluated by adding 10 µg of rPspA or purified PS to the AMP solution prior to incubation with the bacteria. The control group was incubated with 10 µg of bovine serum albumin (BSA) and 25 µg/ml of HNP-1. The number of surviving bacteria was measured in each group after plating and incubation for 16 h at 37 ºC.

### 4.6 Effect of Anti-PspA on pneumococcal killing by HNP-1

To determine the effect of anti-PspA antibodies on pneumococcal killing by HNP-1, the bacteria were opsonized with 10% of anti-rPspA serum for 15 min at 37 ºC prior to treatment with 25 µg/ml of HNP-1 (the serum concentration was based on previous studies performed by the group (30). The control group was pre-incubated with serum from sham immunized mice before treatment. The number of surviving bacteria was measured in each group after plating and incubation for 16 hours.

### 4.7 Choline chloride treatment to remove surface proteins

PspA is anchored to the bacterial cell-wall via choline residues, thus being classified as a choline-binding protein. Chemical removal of choline binding proteins is achieved by washing the bacterial pellets with a 2% choline chloride solution (30). To evaluate the combined effects of polysaccharide capsule and PspA in the action of HNP-1, the capsule-negative mutant strains were grown as described and harvested by centrifugation and divided into two groups. The treatment group was resuspended in 2% choline chloride diluted in PBS and incubated for 10 minutes at room temperature, while the control group was incubated in PBS alone for the same duration. Following incubation, cells were washed twice with PBS to remove residual choline chloride and the eluted proteins, before being immediately used in the bactericidal assay.

### 4.8 Statistical Analysis

All experiments were performed in quadruplicates and repeated to confirm the data. The comparison between wild-type and mutant pneumococcal strains treated with different concentrations of the AMP was performed using two-way ANOVA with Sidak’s post-test. Differences in survival between bacteria treated with recombinant protein and control protein or with control serum and anti-PspA serum were analyzed using Student’s t-test. Statistical significance was calculated based on p≤0.05. All graphs were generated using GraphPad Prism 8.

## 5. ACKNOWLEDGEMENTS

We gratefully acknowledge Dr. David Briles for his invaluable inspiration and foundational contributions to this work, which was motivated by his pioneering efforts in the field and insightful suggestions. His expertise in *Streptococcus pneumoniae* and Pneumococcal surface protein A (PspA), coupled with his generous and kind nature, have profoundly influenced our research.

